# Multifunctionality of V-type ATPase during asexual growth and development of *Plasmodium falciparum*

**DOI:** 10.1101/2023.08.02.551680

**Authors:** Neeta Shadija, Swati Dass, Wei Xu, Hangjun Ke

**Affiliations:** Center for Molecular Parasitology, Department of Microbiology and Immunology, Drexel University College of Medicine, Philadelphia, Pennsylvania, USA, 19129

## Abstract

V-type ATPases are highly conserved hetero-multi-subunit proton pumping machineries found in all eukaryotic organisms. They use ATP hydrolysis to pump protons, acidifying intracellular or extracellular compartments, and are thus crucial for various biological processes. Despite being evolutionarily conserved in malaria parasites, this proton pump remains understudied. To understand the localization and biological function of V-type ATPase in the deadliest human malaria parasite *Plasmodium falciparum*, we utilized CRISPR/Cas9 to endogenously tag the subunit A of the V_1_ domain at the C-terminus. V_1_A (PF3D7_1311900) was tagged with a triple hemagglutinin (3HA) epitope and TetR-DOZI-aptamers for conditional expression under the regulation of anhydrotetracycline. Through immunofluorescence assays, we identified that V-type ATPase was expressed throughout the intraerythrocytic developmental cycle and was mainly localized on the digestive vacuole and plasma membrane. Immuno-electron microscopy further revealed that V-type ATPase was also localized on secretory organelles, such as rhoptries in merozoites. Knockdown of V_1_A led to cytosolic pH imbalance and blockage of hemoglobin digestion in the digestive vacuole, resulting in an arrest of parasite development in the trophozoite stage and, ultimately, parasite demise. Using BN-PAGE/Western blot, we detected a large molecular weight complex (∼ 1.0 MDa) corresponding to the total molecular weights of V_1_ and V_o_ domains. The complex was readily disrupted by the V-type ATPase specific inhibitor Bafilomycin A1, but not by low glucose conditions or treatment with chloroquine. Together, our data suggest that V-type ATPase is localized on several subcellular compartments in *P. falciparum* and plays critical roles to support malaria parasites to grow and replicate inside red blood cells.

## Introduction

Malaria remains one of the deadliest infectious diseases worldwide. In 2021, the WHO reported 247,000,000 malaria cases and 619, 000 deaths^1^. The causative agent of this disease is the protozoan parasite called *Plasmodium*, belonging to the phylum Apicomplexa. Among the five *Plasmodium* species infecting humans (*P. falciparum*, *P. vivax*, *P. ovale*, *P. malariae*, *P. knowlesi*), *P. falciparum* is responsible for the majority of malaria cases and deaths worldwide. All clinical symptoms of malaria arise due to the repetitive growth of the parasites in red blood cells (RBCs); hence, the intraerythrocytic developmental cycle (IDC) or the asexual blood stage has long been recognized as a crucial target for developing antimalarial interventions.

Throughout the IDC, malaria parasites reside in a parasitophorous vacuole and contain many intracellular compartments, including the cytosol, the digestive vacuole (DV) for hemoglobin digestion, the nucleus/ER/Golgi apparatus, the mitochondrion, the apicoplast, and secretory organelles in merozoites. Some of these subcellular compartments are known to have distinct pH, which is key for sustaining the biological processes inside them. For instance, the parasite cytosol maintains a static pH near 7.3^2^, whereas the DV requires a lower pH near 5.0 for hemoglobin digestion^3-6^. The mitochondrion has a slightly basic pH (7.37 ± 0.09) whereas the apicoplast is more neutral (7.12 ± 0.40)^7^.

Regulation of pH in distinct subcellular compartments involves multiple players, including buffers, enzymes, transporters, and proton pumps^8^. Proton pumps are critical because they can directly transport protons against the concentration gradient, actively regulating subcellular pH^9^. In *P. falciparum*, two types of proton pumps are present^10^, including the pyrophosphate (PPi) driven H^+^-pumping pyrophosphatases (H^+^-PPases) or vacuolar pyrophosphatases (V-PPases)^11-13^, and the ATP driven vacuolar type ATPases (V-type ATPases)^14^. VP1 (vacuolar pyrophosphatase 1) is an example of PPi driven proton pumping pyrophosphatase. In *P. falciparum*, a recent study has shown that VP1 is mainly localized on the parasite plasma membrane, regulates cytosolic pH, and plays essential roles in the ring stage and the ring-to-trophozoite transition^15^. On the other hand, V-type ATPase is a large molecular weight ATP hydrolyzing proton pumping machinery^16^, consisting of a V_1_ domain containing subunits of A-H and a V_o_ domain containing subunits of a, c, c’, c’’, d, e. The cytosolic V_1_ domain holds the ATP hydrolysis hexamer made of A and B subunits, whereas the membrane bound V_o_ domain contains the H^+^ channel, which is involved in proton transport across the membrane. A unique feature of this machinery lies in the reversible assembly and disassembly of the V_1_V_o_ domains^17^, regulating the pump’s function in response to various extracellular conditions^18^. The *P. falciparum* genome encodes all V_1_ domain subunits and most V_o_ domain subunits that are present in other eukaryotes (www.PlasmoDB.org). The recent genome-wide mutagenesis study in *P. falciparum*^19^ and the knockout survey in *P. berghei*^20^ reveal that nearly all V-type ATPase subunits are essential in the asexual blood stage. Despite the significance, our understanding of the physiological role of V-type ATPase in malaria parasites remains rather limited.

In classical model organisms, including yeast, humans, and plants, V-Type ATPase is localized on the membranes of multiple subcellular compartments such as endosome, lysosomes, vacuoles, Golgi, and the plasma membrane^21-23^. In marine phytoplankton such as diatoms, dinoflagellates and coccolithophores, V-type ATPase localizes to the membranes of chloroplasts derived from endosymbiosis and significantly enhances carbon fixation and oxygen production^24^. In *P. falciparum*, previous studies have suggested that V-type ATPase is a major proton pump responsible for maintaining cytosolic pH^2^ and DV pH^5^ in trophozoite-stage parasites. However, its localization has remained elusive or somewhat controversial. An early study indicated that V-type ATPase has heterogenous but not well-defined localization^25^. In another study, immunofluorescence assays (IFAs) and immuno-EM suggested that V-type ATPase is localized on the parasite periphery, DV, and small clear vesicles^26^. However, the use of antibodies raised against the subunits of the bovine V-type ATPase in the study^26^ was likely problematic due to limited specificity towards the *Plasmodium* counterparts. Furthermore, another report revealed that V-type ATPase also localizes on the RBC membrane, apparently being exported to the RBC cytosol^27^. Thus, studies of the *Plasmodium falciparum* V-type ATPase have not provided a consensus on its localization. Therefore, further investigation is needed to clarify the localization and function of this important proton pump in malaria parasites.

In this study, we employed genetic and biochemical approaches to better understand the localization and functionality of V-type ATPase in *P. falciparum*. Our data suggest that V-type ATPase is expressed throughout the IDC and exhibits dynamic localization patterns that vary depending on the specific developmental stages of the parasite. We have also discovered the intriguing localization of V-type ATPase on rhoptries and secretory vesicles in merozoites. Moreover, our results underscore the essentiality of V-type ATPase throughout the IDC. Depletion of the proton pump leads to the arrest of trophozoites, which contain undigested hemoglobin in the DV. Using the V-type ATPase specific inhibitor Bafilomycin A1, we have discovered that V-type ATPase potentially plays important roles during parasite egress and invasion. Finally, our study demonstrates that the V-type ATPase is sensitive to disruption by Bafilomycin A1, but the complex stability is less susceptible to certain treatments, such as low glucose or chloroquine.

## Results

### Endogenous tagging of V_1_A via CRISPR/Cas9 for localization and conditional knockdown

The highly conserved V-Type ATPase has 2 domains, the V_1_ domain that hydrolyses ATP and the V_o_ domain carrying out proton translocation using the free energy released from ATP hydrolysis. *P. falciparum* encodes all subunits required to form the V_1_ subunit, including subunits of A (PF3D7_1311900), B (PF3D7_0406100), C (PF3D7_0106100), D (PF3D7_1341900), E (PF3D7_0934500), F (PF3D7_1140100), G (PF3D7_1323200), and H (PF3D7_1306600). The V_o_ includes subunits of a (PF3D7_0806800), c (PF3D7_0519200), c’’ (PF3D7_1354400) and e (PF3D7_0721900). Subunits A and B form the ATP hydrolyzing hexamer to power the rotational movement of the entire proton pump. To study V-type ATPase in *P. falciparum*, we successfully tagged the endogenous locus of subunit A (V_1_A) C-terminally with the triple HA epitope (3xHA) and elements required for conditional knockdown using the TetR-DOZI-aptamer system^28,29^. This was achieved via CRISPR/Cas9^30,31^ with two different gRNA plasmids (**Figure 1A**). We termed the resulted transgenic parasite line NF54attB-V_1_A-3HA^apt^. We also performed limiting dilution to clone the transgenic parasite line to obtain pure parasites. The D11 clone was chosen for all subsequent experiments. The genotype of the NF54attB-V_1_A-3HA^apt^ line (D11 clone) was verified by diagnostic PCR (**Figure 1B**). The expression of V_1_A-3HA proteins at the expected molecular weight was confirmed by Western blot (**Figure 1C**).

**Figure 1.**
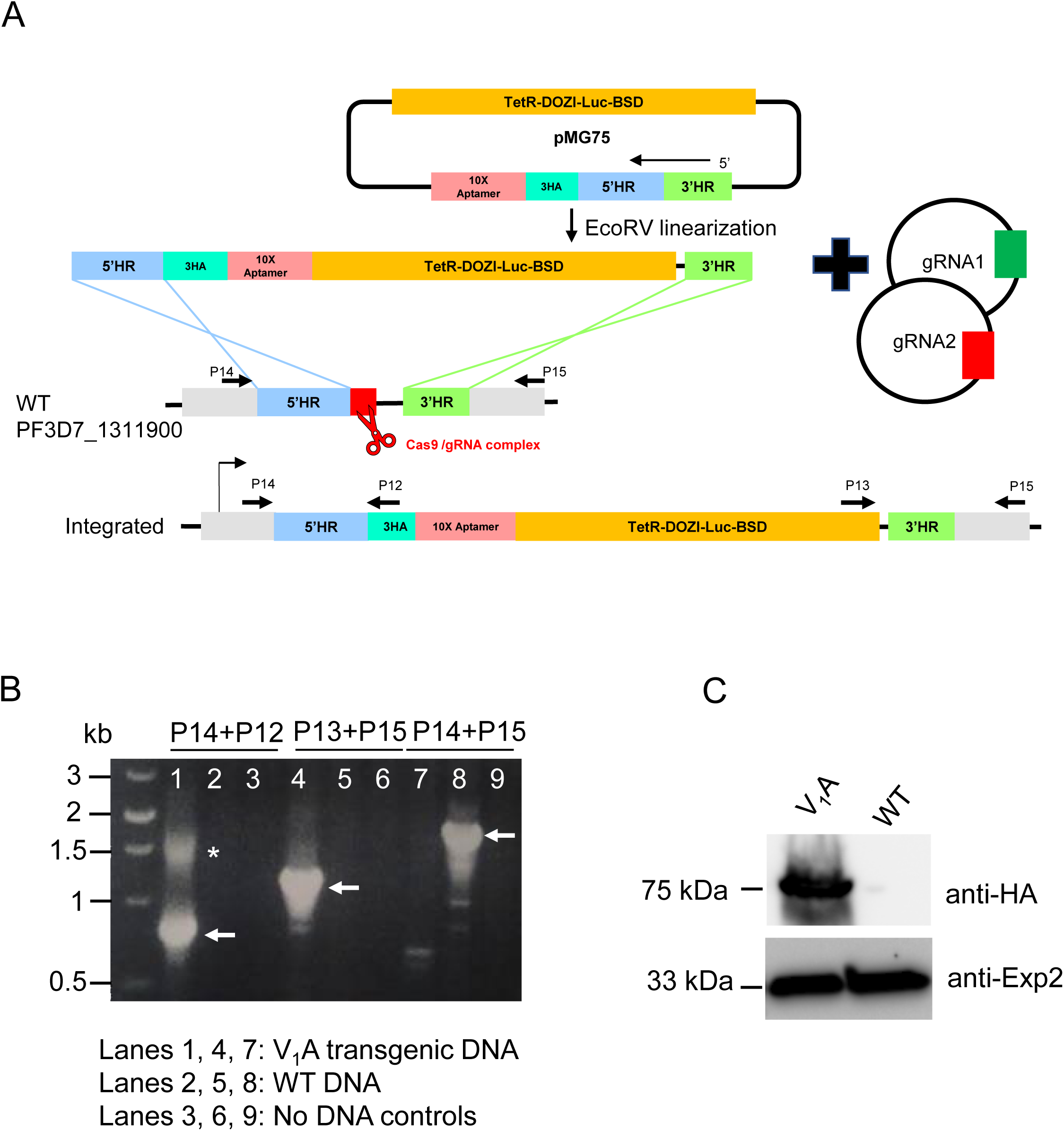
Endogenous tagging of V_1_A via CRISPR/Cas9 for conditional expression. **A**, Schematic of modifying the V_1_A genetic locus via CRISPR/Cas9 mediated double crossover recombination. The primer positions were indicated by black arrows with sequences listed in Table 1. **B**, Diagnostic PCR/DNA electrophoresis showing the corrected modification of the V_1_A genetic locus in the D11 clone of NF54attB-V_1_A-3HA^apt^. The band sizes of PCR products were 0.8 kb (lane 1), 1.2 kb (lane 4), and 1.9 kb (lane 8). One non-specific band was detected in lane 1 (star). **C**, Western blot showing the expression of V_1_A-3HA proteins in the transgenic parasite line, NF54attB-V_1_A-3HA^apt^.

**Table 1.**
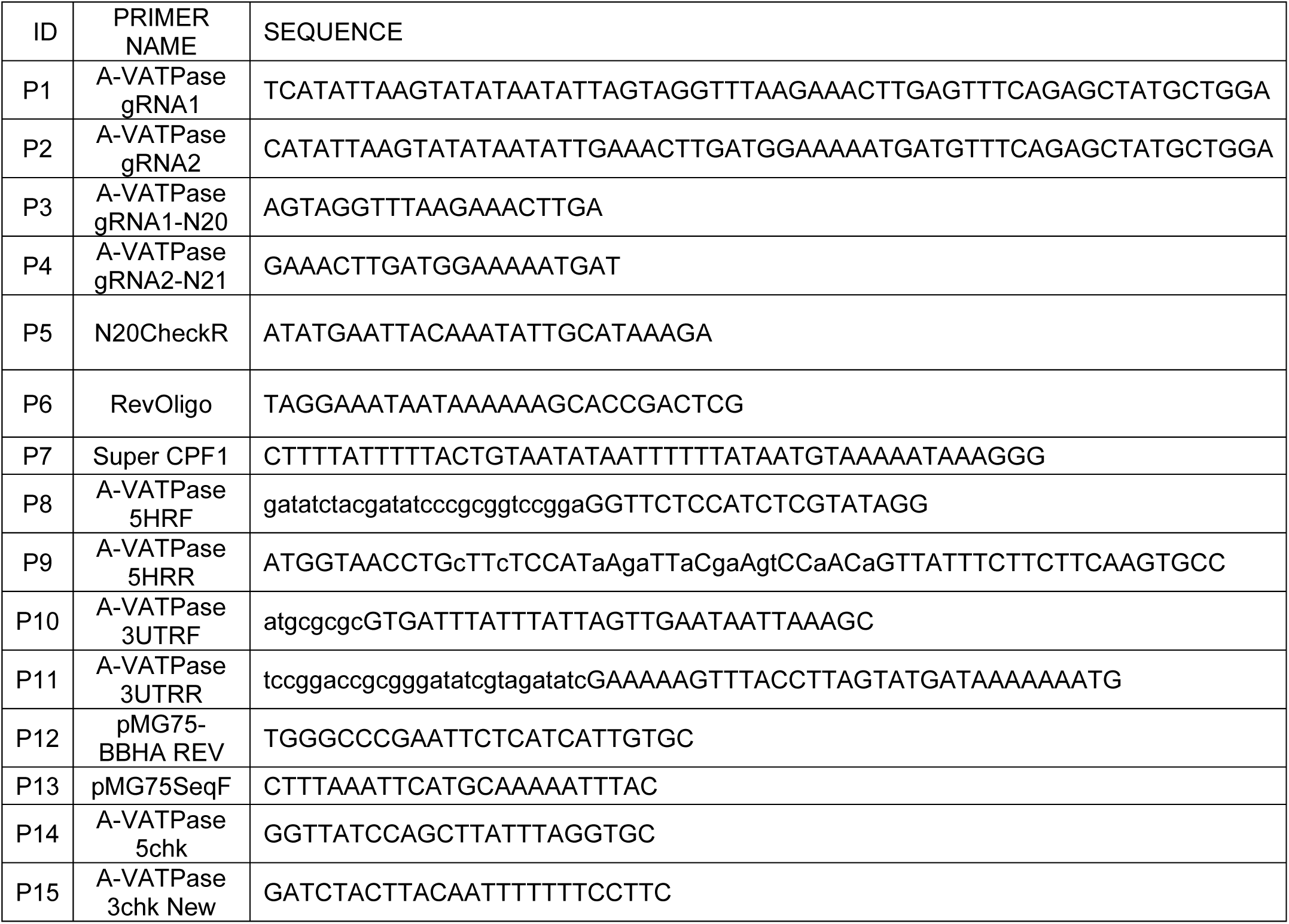
Primers and oligoes used in this study.

To verify the localization of V-type ATPase in *P. falciparum*, we tightly synchronized the parasites with alanine/HEPES and performed Immunofluorescence studies (IFAs) over the 48 h lifecycle. We used Exp2^32^, the maker for the Parasitophorous Vacuolar Membrane (PVM), as the indicator of the parasite periphery. From the time of invasion (T0), we harvested samples every 8 hours in the ring stage and every 4 hours in trophozoite or schizont stages. V_1_A was seen to be expressed in all asexual blood stages (**Figure 2A**). In the ring stage, V_1_A localized to the parasite periphery, showing good colocalization with Exp2. In the trophozoite stage, V_1_A was predominantly localized on both the Parasite Plasma Membrane (PPM) and the Digestive Vacuole (DV). This localization pattern was also observed in the early schizont stage. However, in the late schizont stage, some punctate signals appeared near the parasite periphery. To further characterize this, we performed IFA on schizonts pre-treated with the egress inhibitor, ML10^33^, a reversible blocker of protein kinase G. Treatment of ML10 inhibits parasite egress, leading to an increased number of mature late schizont stage parasites. In the treated mature schizonts, V_1_A’s localization appeared more punctate, lacking confined signals showing circular or membranous structures (**Figure 2B**). This prompted us to further verify V-type ATPase’s localization via Immuno-Electron Microscopy. Indeed, immuno-EM revealed a specific localization of V_1_A on rhoptries and some small vesicles near them (**Figure 3**), indicating that V-type ATPase is also present on the secretory organelles in merozoites. Altogether, we have used IFA and Immuno-EM to uncover the dynamic localization of V-type ATPase in different asexual stages of *P. falciparum*. This dynamic localization of V-type ATPase may be associated with its multifunctionality throughout the 48 h lifecycle.

**Figure 2.**
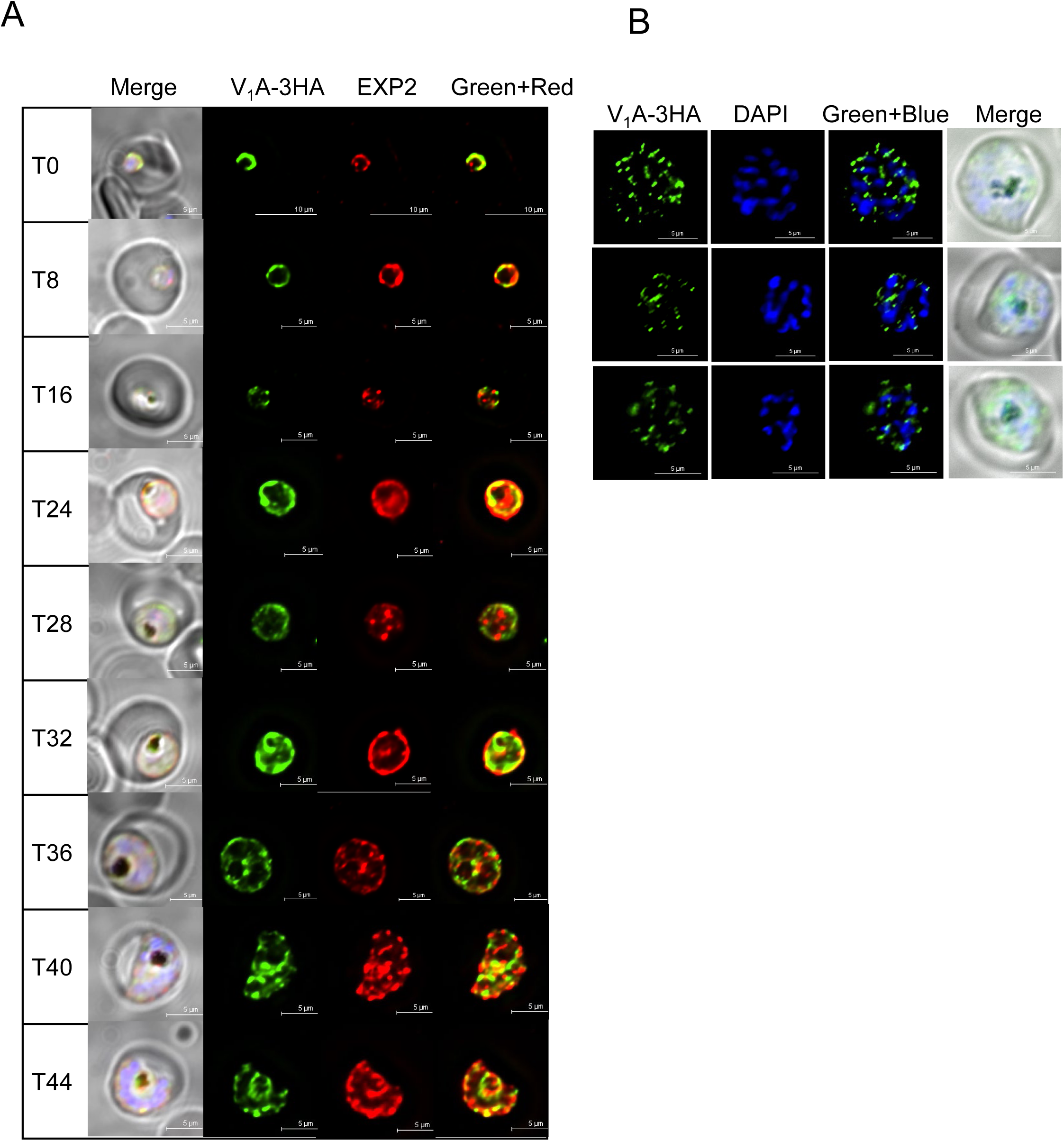
Localization of V_1_A throughout the asexual blood stages. **A**, Localization of V_1_A throughout the lifecycle. In the NF54attB-V_1_A-3HA^apt^ line, V_1_A was detected by anti HA and FITC-labeled secondary antibodies. Exp2 was detected by anti-Exp2^49^ and TRITC-labeled secondary antibodies. **B**, Localization of V_1_A in mature schizont stage parasites treated with ML10. The synchronized late-trophozoite stage parasites were treated with 25 nM of ML10 for 14 h to reach the mature schizont stage. V_1_A was detected by anti HA and FITC-labeled secondary antibodies. DNA was stained with DAPI. In each condition, representative images of at least 25 parasites were shown.

**Figure 3.**
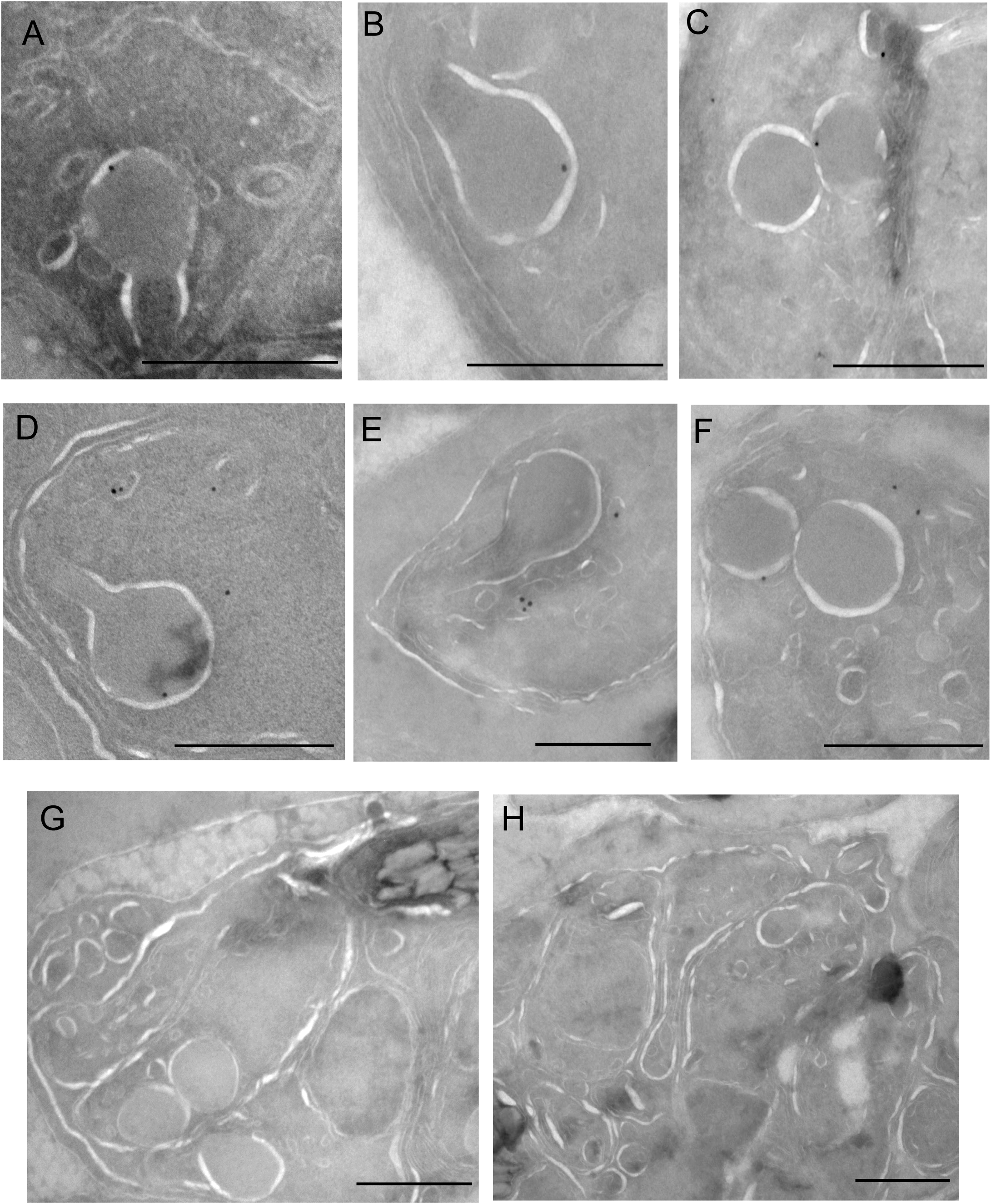
Localization of V_1_A on secretory organelles in mature schizont stage parasites. **A-F**, Detection of V_1_A signals on rhoptries and secretory organelles near rhoptries in the parasite line NF54attB-V_1_A-3HA^apt^ by immuno-electron microscopy. Bars, 500 nm. **G-H**, Negative control images with the primary antibody omitted throughout the experiment. Bars, 500 nm.

### V-type ATPase is essential for asexual growth and development in *P. falciparum*

To understand the essentiality of V-type ATPase in *P. falciparum*, we performed knockdown studies using two different methods to remove aTc. In the first method, we synchronized the parasites several times and started aTc removal in the early ring stage (**Figure 4A**). The knockdown parasites appeared healthy 24 h post aTc removal, but they failed to produce any new ring stage parasites in the next 24 h. At 48 h of knockdown, the parasites were arrested in the trophozoite stage and lysed at 72 h. Thus, V_1_A knockdown from the early ring stage resulted in trophozoite arrestment and parasite demise within one cycle. In the second method, we started aTc removal from the synchronized trophozoite stage parasites (**Figure 4D**). The knockdown parasites were able to produce new ring stage parasites at 24 h post aTc removal, but were arrested again in the trophozoite stage at 48 h. The arrested parasites did not initiate another IDC, but remained arrested or lysed at 72 h. Thus, both knockdown methods revealed the essentiality of V-type ATPase for asexual growth and development in *P. falciparum*. In each experiment, quantification of parasite growth in the aTc (±) conditions was shown in **Figure 4B and 4E**. The V_1_A protein levels of the knockdown experiments were shown by Western blot in **Figure 4C and 4F**.

**Figure 4.**
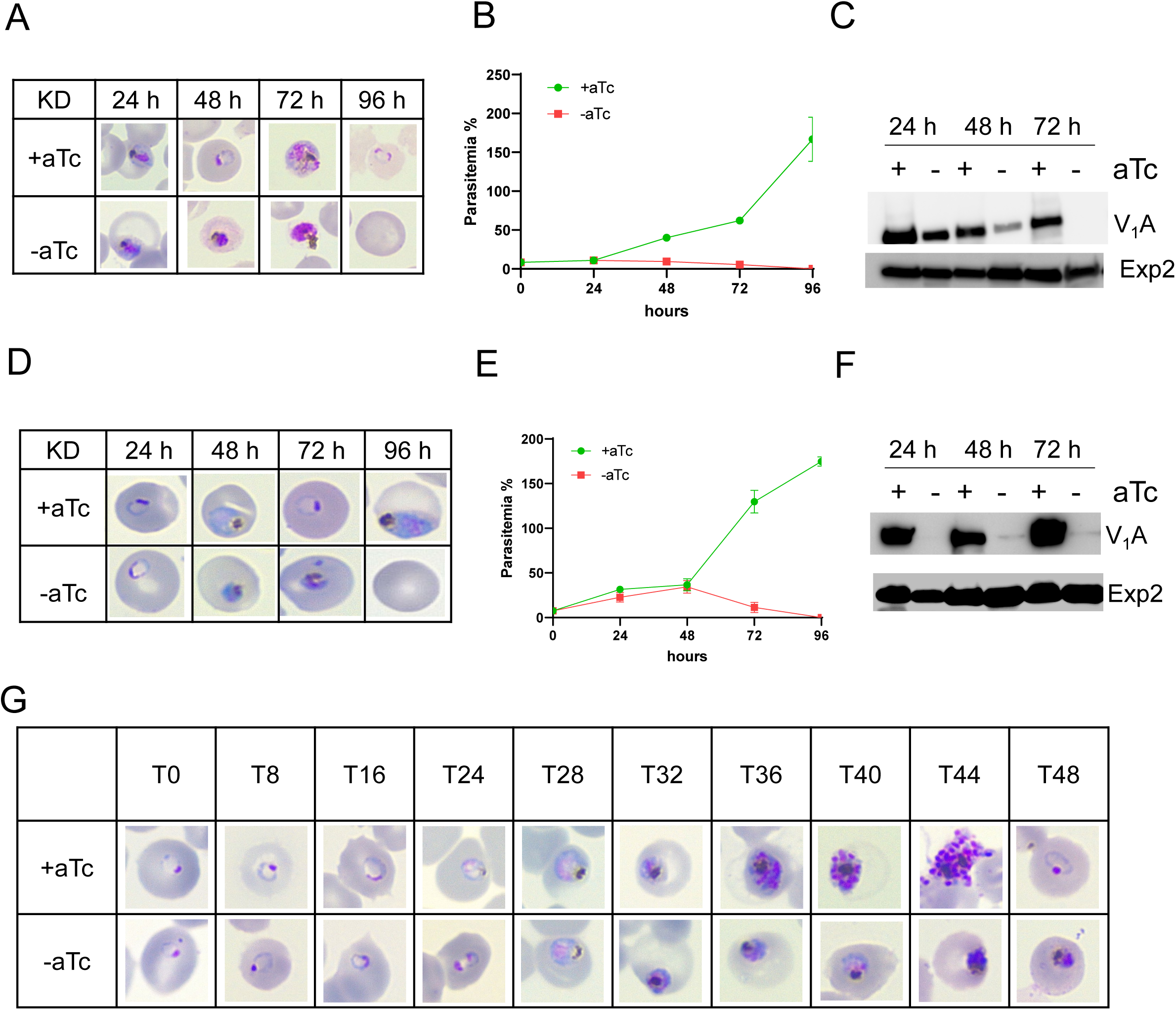
Essentiality of V-type ATPase in the asexual blood stages. Giemsa-stained images showing parasite morphologies resulted from knocking down of V_1_A from the ring stage (**A**), or from the trophozoite stage (**D**), in the NF54attB-V_1_A-3HA^apt^ line. Quantification of parasitemia in the aTc +/- cultures (**B, E**). At each time point, parasitemia was determined from counting 1000 RBCs under a light microscope. Error bars indicated the s.d. of three technical measurements. Western blot showing the expression level of V_1_A in the knockdown experiments (**C, F**). The same blot was re-probed with anti-Exp2 to show loading controls. **G**, Giemsa-stained images showing parasite morphological changes over the time course experiment. Experiments of A-F were repeated three times. Panel G was repeated two times.

To better understand parasite developmental blockage upon V_1_A knockdown, we observed the morphological changes of the parasites in a more detailed time course experiment (**Figure 4G**). In tightly synchronized parasites, we collected thin blood smears every 8 hours during the ring stage and every 4 hours during the trophozoite and schizont stages. Unlike the aTc (+) parasites that completed their development normally, we observed an arrest in parasite growth in the aTc (-) culture at 32 h post aTc removal. The knockdown parasites were arrested in the mid-trophozoite stage and did not progress significantly in the remaining hours of the IDC. Altogether, using different approaches, we have shown that knockdown of V-type ATPase leads to the arrest of parasite development in the trophozoite stage within one IDC post aTc removal. Overall, our results confirm that V-type ATPase is essential for *P. falciparum*, with the trophozoite stage being the most susceptible to the loss of this proton pump.

### V-type ATPase is essential for hemoglobin digestion

V-type ATPase has been shown to regulate the pH of DV in trophozoite stage parasites^3-5^. A low pH (∼ 5.0) is essential for hemoglobin digestion, which is extremely important for malaria parasites to obtain nutrients and gain space to grow inside RBCs^34^. To get a deeper understanding of the knockdown effects on the DV, we performed Transmission Electron Microscopy (TEM) studies. The knockdown experiment was initiated from Percoll-enriched schizont stage parasites, and we purified parasites by a magnetic column at 36 h post aTc removal. The parasitemia of both aTc (±) cultures reached nearly 100% (**Figure 5A**). The aTc (-) culture displayed more cell lysis compared to the aTc (+) culture after magnetic enrichment, indicating that the integrity of some arrested trophozoites was compromised. Under TEM, the knockdown parasites exhibited striking features (**Figure 5B**). The DV of knockdown parasites showed significantly higher electron density than the control. Additionally, we observed membrane-bound large vesicles inside the DV, indistinguishable from the RBC cytosol in terms of electron density. These structures apparently contained large amounts of undigested hemoglobin. Although not measured quantitatively, we also noticed a significant reduction in hemozoin crystals in the knockdown parasites compared to the control. Consequently, knockdown of V-type ATPase leads to defects in several steps of hemoglobin metabolism, including the release of hemoglobin into the DV, breakdown of hemoglobin into small peptides, and formation of hemozoin.

**Figure 5.**
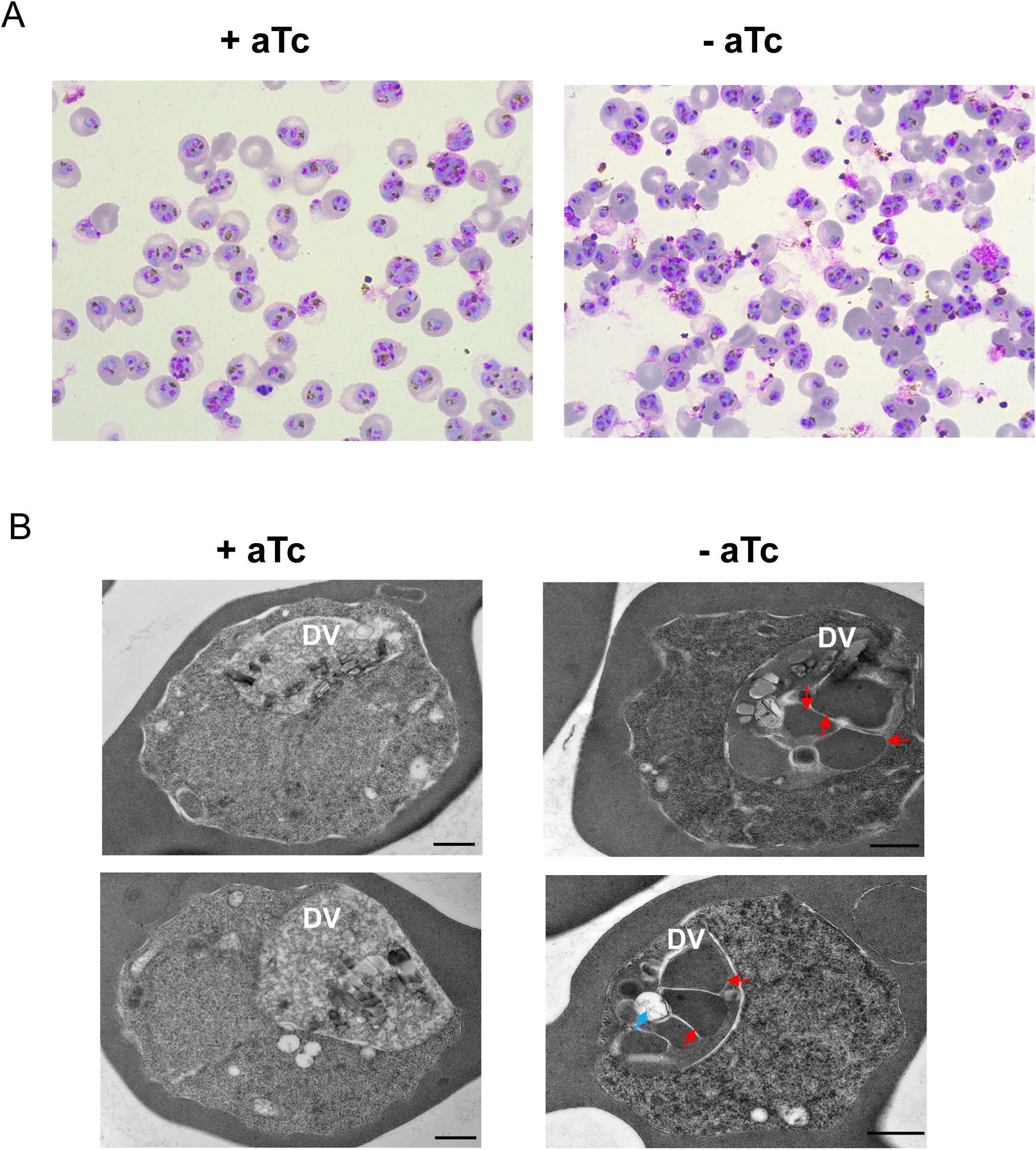
V-type ATPase is essential for hemoglobin digestion in the digestive vacuole. **A**, Giemsa-stained images of control and V_1_A knockdown parasites enriched by a magnetic column. **B**, Electron Transmission Microscopic images of control and V_1_A knockdown parasites. In the aTc - parasites, many vesicles inside the digestive vacuole had electron dense signals and were surrounded by membranes (red arrows). Some vesicles had translucent appearance but were still surrounded by membranes (blue arrow).

Hemoglobin uptake is initiated at a specialized structure called cytostome^35^, and hemoglobin-containing vesicles (HCvs) are pinched off from the membranes and fused with the DV to release their contents for proper digestion and heme detoxification. The hemoglobin containing vesicles are thus wrapped with two membranes, with the PPM on the outside and the PVM on the inside^36^. Interestingly, knockdown of V-type ATPase blocked hemoglobin digestion in the DV, but the upstream trafficking of HCvs enroute to the DV was seemingly not affected. This differed from a previous study that showed accumulation of HCvs inside the parasite cytosol upon conditional inactivation of PfVPS45^37^, a protein involved in vesicular trafficking. Moreover, the fusion of the outer membrane of HCVs to the DV membrane is potentially a rapid process as this event has been rarely observed in wildtype parasites^38^. In the V_1_A knockdown parasites, the undigested hemoglobin was surrounded by a single membrane (**Figure 5B, red arrows**). This information indicated that fusion of HCVs with the DV membrane was likely normal upon depletion of V-type ATPase, but degradation of the inner membrane of HCvs was blocked. Interestingly, we also observed that some vesicles in the DV of the knockdown parasites had much reduced electron density, likely due to a more advanced level of hemoglobin digestion (**Figure 5B, blue arrow**). Overall, our results show that V-type ATPase is critical for hemoglobin release and digestion in the DV.

### V-type ATPase is essential for regulating cytosolic pH

Apart from its role in the DV, V-type ATPase is known to regulate cytosolic pH by pumping protons across the PPM in *P. falciparum*^2^. This understanding was based on the observation that Bafilomycin A1, an inhibitor of V-type ATPase, caused a dramatic decline of cytosolic pH. A recent study suggests that Bafilomycin A1 causes a stearic hinderance and blocks the c-ring rotation by preventing the interaction between subunit a and the c-ring^39^. The effect of V-type ATPase on cytosolic pH, however, has not been confirmed by genetic studies in *Plasmodium*. To address this, we utilized the well-established pH measurement technique with BCECF-AM (2’,7’- Bis-(2-Carboxyethyl)-5-(and-6)-Carboxyfluorescein, Acetoxymethyl Ester) in saponin-treated trophozoites^2^. We quantified cytosolic pH of aTc ± cultures at 36 h post knockdown from Percoll-enriched schizont stage parasites. We included Bafilomycin A1 as a positive control. Our assays revealed that cytosolic pH was 7.24±0.08 in the control parasites but reduced to 6.01±0.28 upon knockdown of V-type ATPase (**Figure 6**). This result confirms that V-type ATPase is the major proton pump for regulating cytosolic pH in trophozoite stage parasites. Consistent with the published data^2^, we also observed that Bafilomycin A1 caused a profound decrease in cytosolic pH from 7.24 to 6.51±0.1 within 5 min of treatment. Interestingly, the combination of V_1_A knockdown and Bafilomycin A1 resulted in an even lower cytosolic pH (5.77±0.15), although this effect was not statistically significant compared to knockdown alone (*p* > 0.05). Overall, our findings confirm that V-type ATPase plays a major role in regulating cytosolic pH in the trophozoite stage parasites.

**Figure 6.**
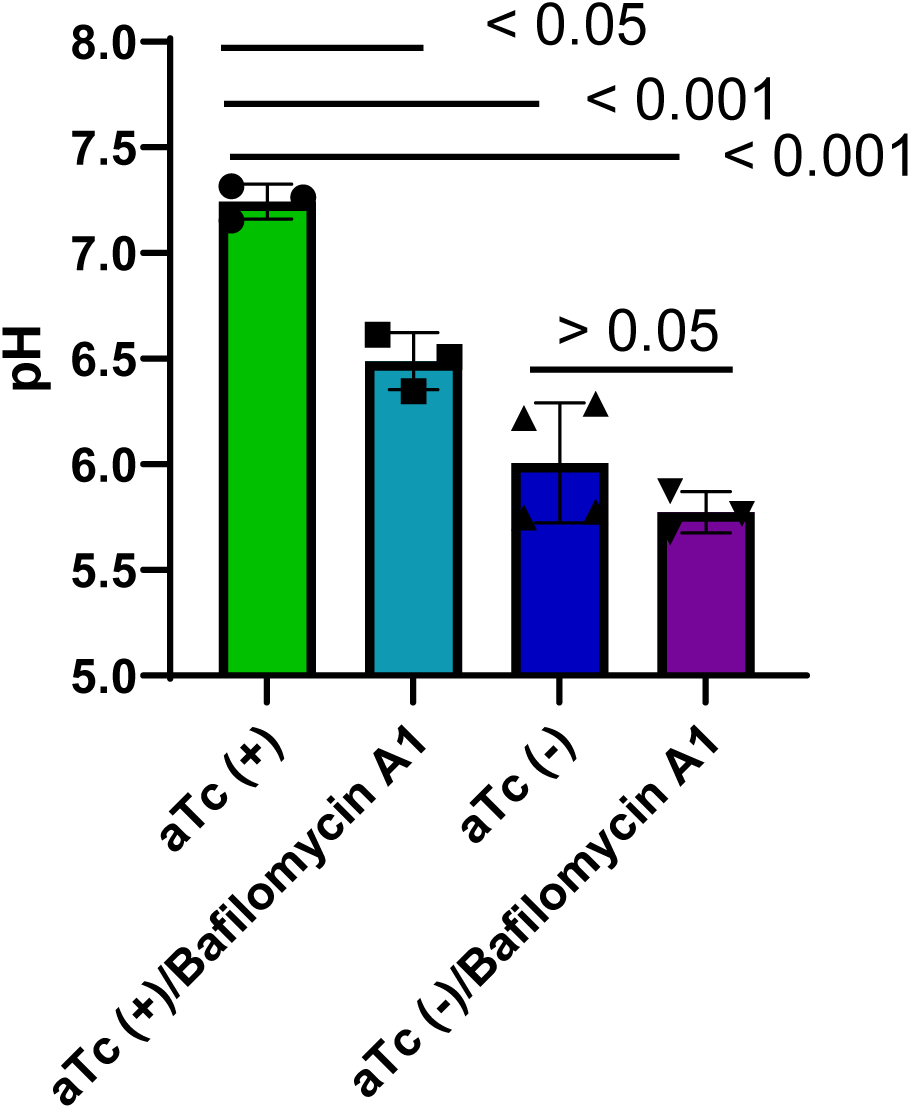
V-type ATPase regulates cytosolic pH in trophozoite stage parasites. At 36 h post knockdown, aTc +/- cultures of the NF54attB-V_1_A-3HA^apt^ line were subjected to pH measurement using the BCECF-AM method in the absence of presence of 100 nM Bafilomycin A1 (100 nM) for 5 min. Error bars indicated the s.d. of 3-4 measurements in each condition. Statistical analysis was done by student t-test. This experiment was repeated two times.

### Viability of the V-type ATPase knockdown parasites

Our data have shown that knockdown of V-type ATPase led to the arrest of trophozoite-like parasites that contained large amounts of undigested hemoglobin in the DV and reduced cytosolic pH. To understand the viability of these parasites, we performed an aTc addback experiment post knockdown (**Figure 7A**). We initiated the knockdown from tightly synchronized Percoll-enriched schizont stage parasites, and performed the addback at different time points post knockdown, 28 h, 36 h, 40 h, and 48 h. Consistent with **Figure 4G**, the knockdown parasites at 28 h did not display obvious morphological defects in Giemsa-stained smears (**Figure 7B**). Upon aTc addback, these parasites were able to produce new ring stage parasites in the next 24 h. At 36 h post knockdown, a delay in parasite development was readily detectable. Upon addback, however, most of the arrested parasites managed to produce new ring stage parasites in the next 24 h. These ring stage parasites continued to become trophozoites at 48 h of addback. This result suggested that the arrested parasites at 36 h post knockdown remained largely viable, despite having severe defects in hemoglobin digestion and pH regulation. At 40 h post knockdown, a smaller percentage of the arrested parasites still succeeded in producing new ring stage parasites at 24 h post addback. However, just a few hours later at 48 h post knockdown, aTc addback failed to rescue any parasite growth. The knockdown parasites either displayed abnormal morphologies or lysed. Altogether, these results imply that *P. falciparum* parasites can sustain viability in a severely stressed condition for at least several hours. Remarkably, aTc addback allowed the arrested parasites to recover from various defects caused by V_1_A knockdown, such as reduced cytosolic pH, blockage of hemoglobin digestion in the DV, and other possible disruptions.

**Figure 7.**
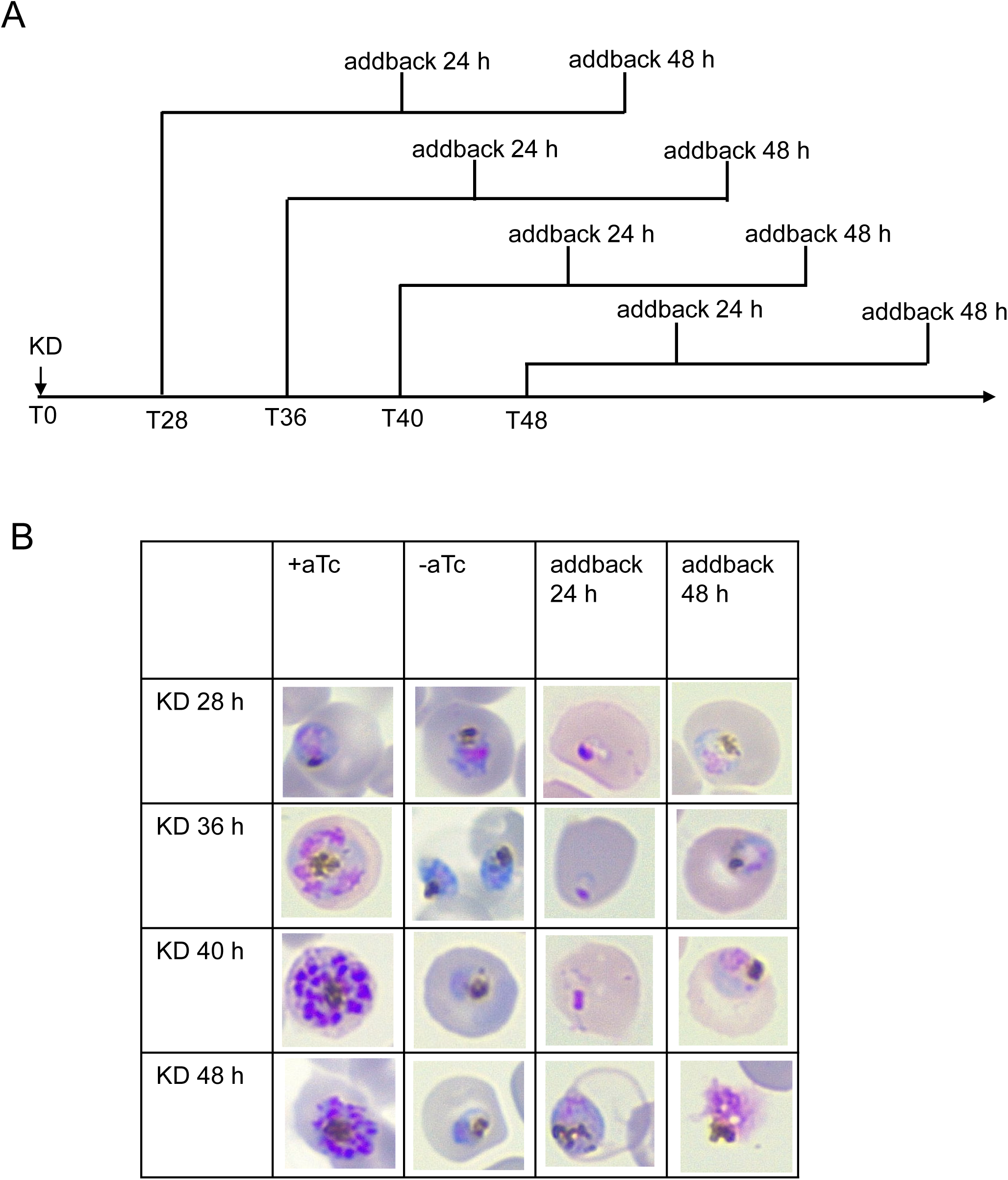
Viability of the V-type ATPase knockdown parasites. **A**, Schematic diagram of the knockdown/addback experiment. At various time points post knockdown, the cultures were fed with aTc + medium and further monitored for 48 h. **B**, Giemsa-stained images showing morphological changes of the parasites upon aTc addback. This experiment was repeated three times.

### The effect of Bafilomycin A1 on parasite egress and invasion

The localization of V-type ATPase on rhoptries and other secretory organelles prompted us to investigate the potential role of this proton pump in parasite egress and invasion. However, in our knockdown studies initiated from the ring, trophozoite, or schizont stage (**Figures 4-7**), an extended period of ∼ 30 h was consistently required before the onset of detectable defects. In addition, all knockdown approaches led to the arrest of trophozoites, making examinations of schizonts infeasible. To address the delayed effects of genetic knockdown, we reasoned that a pool of pre-existing V_1_A proteins, present before aTc removal, might permit parasite survival and maturation over an extended period until the proteins were fully exhausted. To overcome this limitation, we used Bafilomycin A1 to rapidly block V-type ATPase in the late schizonts and observed its effect on parasite egress and invasion. As shown previously, Bafilomycin A1 reduced cytosolic pH from ∼ 7.3 to ∼ 6.5 in about 5 min in *in vitro* assays (**Figure 6**).

In brief, we tightly synchronized the parasites with alanine/HEPES and used ML10 to block parasite egress, thereby increasing the percentage of mature schizonts. At the mature schizont stage (∼ 0 h), we added Bafilomycin A1 (100 nM) for 2 h, washed it out, and allowed parasites to invade new RBCs for 4 h. At 6 h post the initiation of the experiment, we counted the parasitemia of newly formed ring stage parasites and used it as the readout of successful egress and invasion (**Figure 8A**). Parasite growth and morphology were also examined at 24 h and 48 h post the initiation of the experiment. At 6 h, while the new ring stage reached 4.77%±0.25% in the control culture, the Bafilomycin A1 treated culture had a significantly lower parasitemia (0.24%±0.07%) (**Figure 8B**). In the treated culture, events of merozoite invasion were commonly observed, however, formation of new rings was drastically reduced (**Figure 8C**). In the next 24 h and 48 h, the control culture followed the normal progression of the lifecycle. In contrast, the treated culture barely increased parasitemia (**Figure 8B**). The parasites displayed abnormal morphologies or failed to invade RBCs. It was interesting to note that in the treated culture, some merozoite-like parasites appeared to be attached to the RBC surface even at 48 h post the initiation of the experiment (**Figure 8C**). Collectively, treatment of Bafilomycin A1 in the mature schizont stage leads to dramatic reduction in the formation of new ring stage parasites.

**Figure 8.**
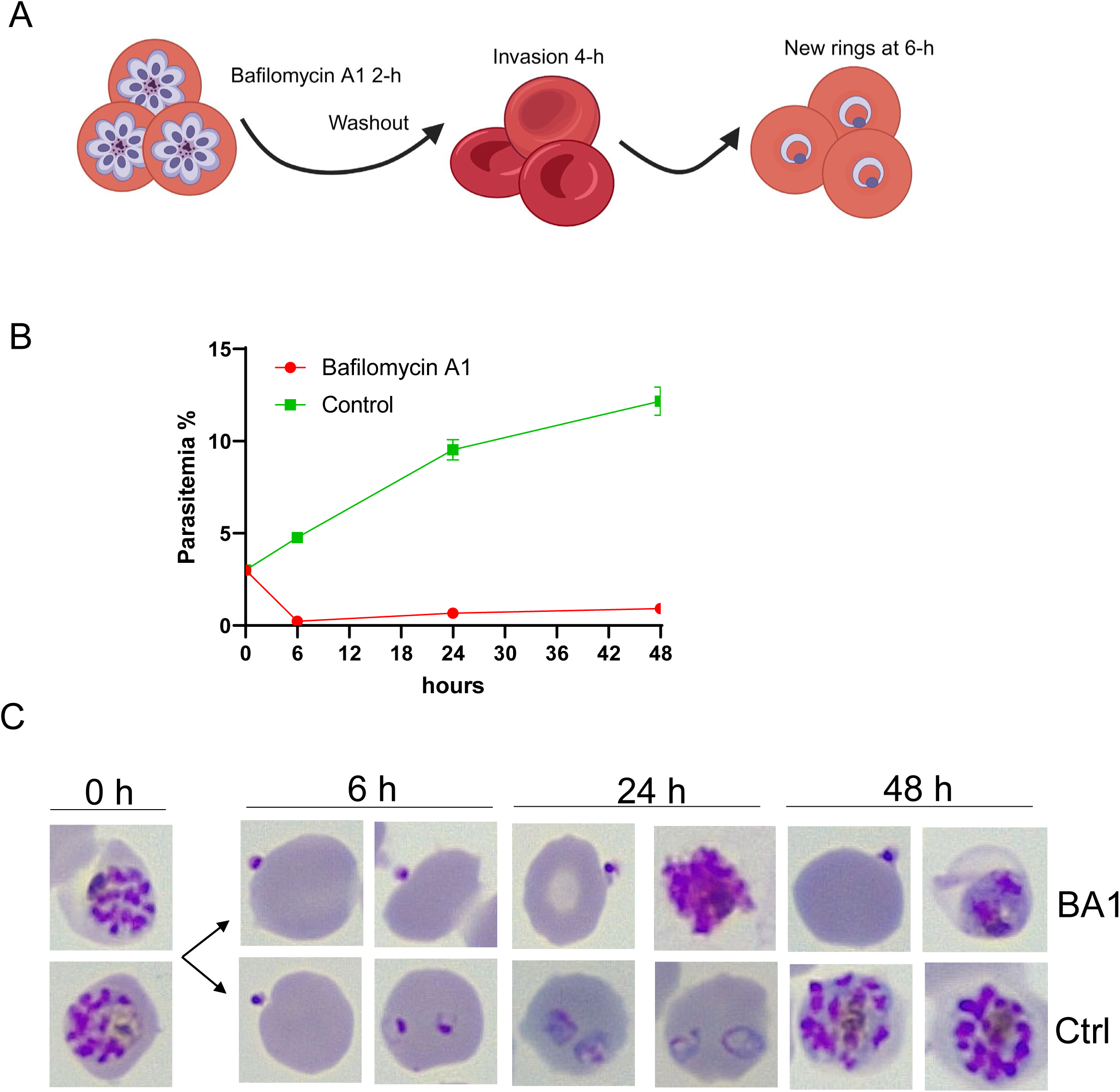
Effects of Bafilomycin A1 on parasite egress and invasion. **A**, Schematic illustration of the experimental design generated by BioRender. Mature schizont stage parasites (NF54attB-V_1_A-3HA^apt^ under 250 nM) were treated with Bafilomycin A1 (100 nM) for 2 h. The drug was washed out and parasites were allowed to invade new RBCs for 4 h. At 6 h, the parasitemia of newly formed ring stage parasites was monitored. Parasite morphology and number were further examined at 24 h and 4 h post the initiation of the experiment. aTc (250nM) was maintained throughout the experiment. **B**, Giemsa-stained imaging showing parasite morphologies in the control and treated cultures. **C**, Quantification of parasite growth throughout the experiment. Parasitemia was determined by counting 1000 RBCs in each smear under a light microscope. Error bars indicated the s.d. of 3 measurements at each time point. This experiment was repeated five times.

### Stability of the V-type ATPase complex under various conditions

In model eukaryotes, it has been well established that V-type ATPase undergoes a reversible process of assembly and disassembly to rapidly regulate its function upon encountering different conditions^17^. For example, the yeast V-type ATPase is very sensitive to the bioenergetic status of the cell and V_1_ and V_o_ disassociate in a low glucose condition^40^. Disassociation of the two domains completely abolishes the proton pumping function as well as the ATP hydrolysis activity of the V_1_ domain^41^. To assess if the *Plasmodium falciparum* V-type ATPase can regulate its function by changing complex stability, we treated synchronized trophozoite stage parasites with Bafilomycin A1, chloroquine, and 1 mM glucose and verified V_1_V_o_ complex formation through BN/PAGE. In parallel, the expression level of the V_1_A monomers was monitored by SDS/PAGE. As shown in **Figure 9**, in the control protein lysate, we detected a high-molecular-weight band near the marker of 1.026 MDa, which approximately corresponded to the total molecular weights of the entire V-type ATPase^21^. In addition, in BN/PAGE, smaller bands near the markers of 720 kDa, 240 kDa, and 66 kDa were also detected. The nature of these smaller complexes is uncertain at present. They could be assembly intermediates of the V_1_V_o_ complex or electrophoretic degradation products, although enough protease inhibitor cocktails were included throughout the experiment. Upon Bafilomycin A1 (10 nM) treatment, the top band near 1.026 MDa was detectable after 2 h, but gradually decreased in intensity after 4 h and 8 h. By contrast, the monomer V_1_A remained more stable throughout the time course. This result indicated that the V-type ATPase dissembled upon Bafilomycin A1 treatment longer than 2 h. Upon treatment with chloroquine (100 nM) or 1 mM glucose, however, the top band near 1.026 MDa remained largely constant even after 8 h. The V_1_A monomer was stably expressed in chloroquine treated samples and gradually decreased in the low glucose treated samples. Together, these data implied that the V-type ATPase complex was readily disrupted by Bafilomycin A1, but not by treatment of chloroquine or a low glucose condition.

**Figure 9.**
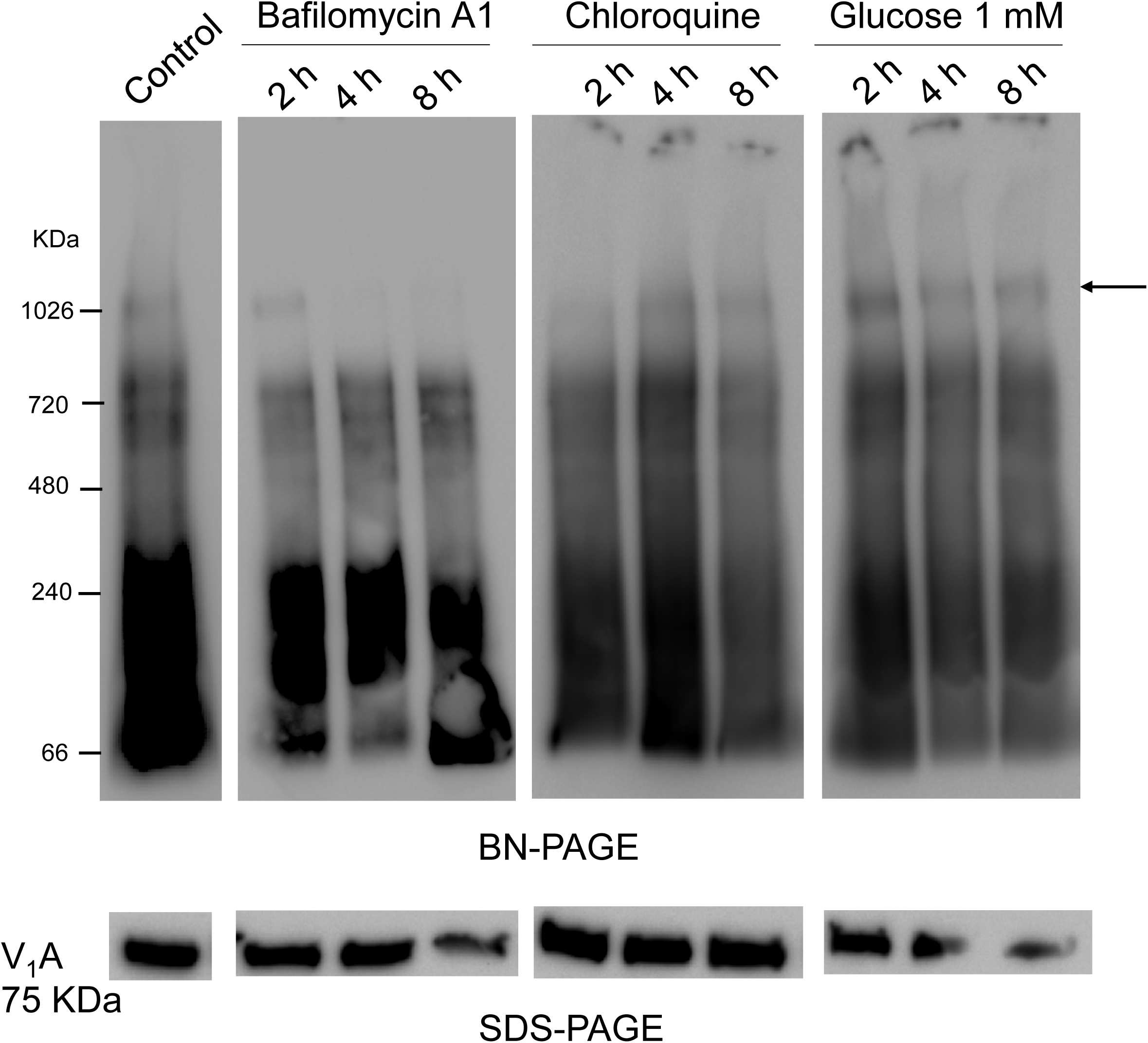
The stability of V-type ATPase complex under various conditions. Synchronized trophozoite stage parasites (NF54attB-V_1_A-3HA^apt^ under 250 nM) were treated with Bafilomycin A1 (10 nM), Chloroquine (100 nM), or 1 mM glucose for 2 h, 4 h, and 8 h. Samples were harvested and analyzed by BN-PAGE and SDS-PAGE. The arrow indicated the large molecular weight band corresponding to the estimated size of the *P. falciparum* V-type ATPase. aTc (250nM) was maintained throughout the experiment.

## Discussion

In this study, we have characterized the essentiality, localization, and biological function of V-type ATPase in *Plasmodium falciparum*. By tagging the subunit A of V_1_ via CRISPR/Cas9, we generated a conditional knockdown parasite line under aTc regulation and obtained several interesting results.

We have found that V-type ATPase is essential for *P. falciparum* during asexual growth and development, with the trophozoite stage being particularly vulnerable to death when this proton pump is depleted (**Figures 4-7**). The localization of *P. falciparum* V-type ATPase is highly dynamic throughout the 48 h asexual lifecycle (**Figures 2-3**). Specifically, in trophozoite stage parasites, V-type ATPase localizes on both the parasite plasma membrane (PPM) and digestive vacuole (DV). This localization pattern is consistent with its role in regulating cytosolic pH^2^ and DV pH^5^. We also observed a drastic reduction in cytosolic pH in the V_1_A knockdown parasites during the trophozoite stage (**Figure 6**). Moreover, our data suggest that V-type ATPase is also expressed and localized on the PPM in the ring stage. However, we need to be cautious when defining the precise localization of V-type ATPase in the small ring stage parasites. A recent study implied that V-type ATPase might not play a major role in proton pumping during the ring stage, as the pyrophosphate driven proton pump, PfVP1, is essential for this stage^15^. Conditional depletion of PfVP1 blocks ring stage development and the ring-to-trophozoite transition, which is apparently not rescued by the V-type ATPase. This indicates that V-type ATPase might either not fulfill proton pumping across the PPM in the ring stage or have a negligible role. In the late schizont stage, we have observed punctate signals of V_1_A in IFA without defined membranous staining (**Figure 2B**). Immuno-EM further revealed V_1_A’s localization on rhoptries and secretory organelles near rhoptries (**Figure 3**). To our best knowledge, this is the first report that observed such localization of V-type ATPase on mature secretory organelles in protozoa. A previous study highlighted the function of V-type ATPase in facilitating the maturation of secretory organelles in *Toxoplasma gondii*, an apicomplexan parasite related to *Plasmodium*^42^, but the presence of V-type ATPase on mature rhoptries or micronemes was not detected^42^. Thus, the unique localization of V-type ATPase on rhoptries and secretory organelles in *Plasmodium* has led us to investigate the role of this proton pump in parasite egress and invasion. To alleviate the slow effect of genetic knockdown, we used Bafilomycin A1 to chemically inhibit V-type ATPase in a short time (2 h) and observed serious defects of the parasites in initiating the next lifecycle (**Figure 8**). Bafilomycin A1’s specificity as a V-type ATPase blocker has been well documented and it has been used as a standard reagent in *P. falciparum* studies^2,5^. We quantified the number of new ring stage parasites post Bafilomycin A1 treatment and used it as a readout to indicate the overall effects on parasite egress and invasion. However, our data did not provide detailed mechanisms underlying the mode of action of Bafilomycin A1 in mature schizont stage parasites. It is possible that Bafilomycin A1 exerts its effects through multiple mechanisms, such as killing the schizonts, blocking egress, and inhibiting invasion. Future investigations will be necessary to characterize these processes in detail. Nevertheless, our data implied that V-type ATPase likely plays a role in parasite egress and invasion.

In the trophozoite stage, we have discovered that V-type ATPase is extremely important for hemoglobin digestion in the DV. This finding aligns with the previously reported role of V-type ATPase in maintaining DV pH^5^. However, observation of undigested hemoglobin in the DV of V_1_A knockdown parasites was astonishing and stands out as a novel finding of this study (**Figure 5**). To date, the processes of hemoglobin uptake from cytostomes and the delivery of hemoglobin containing vesicles (HCvs) to the DV remain enigmatic in *P. falciparum,* as these processes are likely dynamic, transient, and difficult to capture. A previous study reported the rare fusion of one HCv with the DV membrane among > 600 examined thin sections under transmission electron microscopy^38^. Since cytostomes are invaginated from the PPM and PVM, the HCvs are enclosed by the PPM on the outside and the PVM on the inside. The fusion event indicates that the PPM of HCvs fuses with the DV membrane^38^. Therefore, the membrane surrounding the undigested hemoglobin in our data should be the PVM, not the PPM. We can speculate that after HCvs are fused with the DV membrane, the leftover PVM needs to be digested first by specific enzymes (likely lipases) before hemoglobin can be digested by aspartic proteases and cysteine proteases^43^. The enzyme responsible for digesting the PVM of HCvs remains unknown, but it is evidently pH dependent and not among the list of enzymes known to digest hemoglobin^44^.

Moreover, we have detected the native *P. falciparum* V-type ATPase complex by BN/PAGE. In addition to the large molecular weight band near 1.0 MDa, many smaller subcomplex-like moieties were also detected at various molecular weights (**Figure 9**). Though unknown, we speculate that formation of the *P. falciparum* V-type ATPase complex is a highly regulated process, and the intermediate complexes might be involved in the assembly and/or disassembly processes. In mammalian cells, the V-type ATPase was also examined by BN/PAGE but few intermediate complexes were detected^45^. In addition, we noted that the complex was readily disrupted upon Bafilomycin A1 treatment but remained more stable upon challenges of chloroquine and low glucose. The glucose effect on *P. falciparum* V-type ATPase was not expected since low glucose levels are known to dissociate the complex in yeast^40^. This could reflect the unique features of V-type ATPase in *P. falciparum* in a bioenergetically different environment from that of yeast.

In summary, our data have provided valuable insights into the localization and functions of V-type ATPase in *Plasmodium falciparum*, highlighting its critical roles in regulating pH in various subcellular compartments. As we prepared this manuscript, a very recent report by Alder and colleagues also unveiled the localization and function of the *P. falciparum* V-type ATPase, using an inducible knockout approach^46^. Their study has characterized defects related to knockdown of subunit B in the V_1_ domain and subunits a and c in the V_o_ domain. Interestingly, our findings from knockdown of V_1_A align well with those obtained from inducible knockout of V_1_B. Given that subunits A and B work together in the ATP hydrolysis center, genetic knockdown (or knockout) of either A or B leads to common downstream effects. Nonetheless, the combined studies on the two major subunits of V-type ATPase, both from our work and others^46^, have collectively revealed the dynamic localization of V-type ATPase in the asexual blood stage of *P. falciparum*. These studies shed light on the significant role of V-type ATPase in parasite growth and replication, thereby enhancing our understanding of *Plasmodium* biology. Moreover, results of our work and Alder *et al.*^46^ could provide valuable guidance for drug development strategies as V-type ATPase was implicated as a drug target^47,48^.

## Materials and Methods

### 1, Plasmid Construction

Subunit A of *Plasmodium falciparum* V-Type ATPase (V_1_A, PF3D7_1311900) was endogenously tagged with a 3xHA epitope and the TetR-DOZI-Aptamer system at the C-terminus via CRISPR/Cas9. Two NFCas9/gRNA constructs were made according to our cloning procedures published earlier^50^. The DNA oligoes used for cloning and sequencing were listed in **Table 1** (P1-P7). To make the pMG75 template plasmid for double crossover recombination, the 5’ and 3’ homologous regions (5’HR and 3’HR) of V_1_A were PCR amplified from the wildtype genomic DNA using primers P8 and P9 for 5’HR and P10 and P11 for 3’HR. The PCR product was purified and then digested with SacII/BstEII for 5’HR and BssHII/SacII for 3’HR. The digested fragments were sequentially cloned into the pMG75 vector bearing Blasticidin deaminase by T4 DNA ligase. The clones were screened by proper restriction digestion. The positive clones were further confirmed by Sanger sequencing using the primers P12-P13. The final maxi-prep plasmid of pMG75-BSD-V_1_A-3HA was linearized with EcoRV before being transfected into parasites.

### 2, Parasite Culture and Transfection

*Plasmodium falciparum* wildtype strain NF54attB was generously provided by Dr. Joshua Beck (Iowa State University). This line is compatible with all three transfection markers that are commonly used, including BSD (Blasticidin deaminase), hDHFR (human dihydrofolate reductase) and yDHODH (yeast dihydroorotate dehydrogenase). *P. falciparum* was cultured in human O^+^ red blood cells (5% hematocrit, Interstate Blood Bank, Inc.) in the standard RPMI 1640 medium containing 0.5% AlbuMAX® II (Invitrogen), 15 mM HEPES, 10 mg/L hypoxanthine, 25 mM NaHCO_3_, and 50 mg/L gentamicin. Transfections were performed using the standard methods in the ring stage with details shown previously^50,51^. The concentration of aTc was 250 nM in all aTc (+) cultures.

### 3, Immunofluorescence assays (IFA)

Parasites were tightly synchronized with alanine (0.5 M)/HEPES (10 mM) several times. At each time point of the experiment, ∼ 50 µL parasitized RBCs were collected and fixed with 4% paraformaldehyde, 0.0075% glutaraldehyde in 1X PBS. Blocking was done by 3% BSA/PBS for overnight at 4°C. Fixed cells were permeabilized by Triton X-100 (0.25%)/PBS and incubated with primary antibodies overnight at 4°C, including α-HA (1:300, mouse, sc-7392, Santa Cruz Biotechnology) and α-Exp2 (1:500, rabbit)^49^. Following thorough washing, the samples were incubated with FITC- or TRITC- labeled secondary antibodies for 3 h at room temperature. Other procedures followed standard IFA protocols. Images were captured using the Nikon Ti microscope.

### 4, Western Blot for BN-PAGE and SDS-PAGE

Parasite cultures were lysed by Saponin (0.05%)/PBS and host hemoglobin was removed by several washes with 1X PBS. From one culture condition, two aliquots of samples were obtained for BN-PAGE and SDS-PAGE, respectively. For BN-PAGE, the saponin lysed pellet was solubilized with 1% glycodiosgenin (GDN) in MESH buffer (250 mM sucrose, 10 mM HEPES, 1 mM EDTA, 1X protease inhibitor cocktail) for overnight at 4°C. The supernatant after a high-speed centrifugation (13,000 rpm, 10 min) was used for Blue-Native gel electrophoresis and Western blot. For SDS-PAGE, the saponin lysed pellet was solubilized by 2% SDS/Tris (65 mM, pH 6.8) for overnight at 4°C. The supernatant after a high-speed centrifugation (13,000 rpm, 10 min) was used for denatured electrophoresis. For all samples, the protein concentrations were determined by the Pierce™ BCA Protein Assay Kit (ThermoFisher Scientific) to ensure equal loading. Antibodies used for Western blot included α-HA (1:10,000, mouse, sc-7392, Santa Cruz Biotechnology), α-Exp2 (1:10,000, rabbit)^49^, HRP-conjugated goat anti-mouse secondary (1:10,000, A16078, ThermoFisher Scientific), HRP-conjugated goat anti-rabbit secondary (1:10,000, 31460, ThermoFisher Scientific). The membranes were incubated with Pierce ECL Western Blot Substrates and developed using the Bio-Rad chemidoc imaging system.

### 5, Transmission Electron Microscopy

Parasites were synchronized with alanine (0.5 M)/HEPES (10 mM) several times and the schizont stage parasites were enriched by Percoll. After several washes with 1X PBS, the schizonts were added with new RBCs and cultured in aTc +/- conditions for 36 h. Both cultures were then passed through a MACS magnetic column (MiltenyiBiotec) to enrich trophozoite stage parasites, which were fixed with freshly made 2% paraformaldehyde, 2.5% glutaraldehyde in 100mM sodium cacodylate buffer for overnight at 4°C. The samples were shipped to Dr. Wandy Beatty from the Molecular Microbiology Imaging Facility at Washington University, St. Louis, MO for further processing and imaging. Other procedures were carried out as shown previously^51^.

### 6, Immuno-Electron Microscopy

Parasites were synchronized with alanine (0.5 M)/HEPES (10 mM) several times and the trophozoite stage parasites were enriched using a MACS magnetic column (MiltenyiBiotec). The enriched parasites were treated with 25 nM ML10^33^ for 14 h and then fixed with freshly made 4% paraformaldehyde/0.05% glutaraldehyde/100mM PIPES buffer for overnight at 4°C. The samples were shipped to Dr. Wandy Beatty from the Molecular Microbiology Imaging Facility at Washington University, St. Louis, MO for further processing and imaging. Other procedures were carried out as previously published^50^. Antibodies used in the study included α-HA (1:300, mouse, sc-7392, Santa Cruz Biotechnology) and goat anti-mouse IgG 18 nm colloidal gold-conjugated secondary antibody (1:100, Jackson ImmunoResearch Laboratories). A parallel sample with the primary antibody omitted was used as the negative control, which did not react to the secondary antibody.

### 7, pH measurement with BCECF-AM, a ratio-metric pH indicator

pH measurement was carried out according to the published protocols^2^. Post 36 h of aTc removal, the control and knockdown cultures were incubated with 4 µM BCECF-AM (2’,7’-Bis- (2-Carboxyethyl)-5-(and-6)-Carboxyfluorescein, Acetoxymethyl Ester, Fisher Scientific) for 30 min at 37°C. Pluronic at 1:1000 dilution was added to increase cellular permeability to BCECF-AM. The parasites were treated with rapid saponin lysis to remove hemoglobin. The pellet was washed two times with saline/glucose buffer (NaCl 125 mM, KCl 5 mM, MgCl2 1 mM, glucose 20 mM, HEPES 25 mM, pH 7.4) and resuspended in 1 mL of warm saline/glucose buffer. The sample’s fluorescence was monitored using a fluorescence spectrometer (Hitachi F-7000) at 535 nm with excitation wavelengths set at 490/440 nm. When Bafilomycin A1 (100 nM) was used in aTc +/- conditions, the drug was incubated with saponin-lysed and washed parasites for 5 min before they were further analyzed by Hitachi F-7000. The fluorescence ratios were back calculated to pH values according to a linear regression curve generated from several known pH standards (pH 6.8, 7.1, 7.4), as shown previously^2^.

## Acknowledgement

We are grateful for the technical assistance and intellectual input from members of the Center for Parasitology at Drexel University College of Medicine. We thank Dr. Joshua Beck (Iowa State University) for providing the parasite line, Drs. Jaquin Niles (MIT) and Sean Prigge (Johns Hopkins University) for providing the TetR-DOZI-aptamer plasmids, Drs. Baker and Ooij for providing the compound, ML10. We thank Dr. Wandy Beatty at Washington University in St. Louis for performing transmission electron microscopy and immuno-electron microscopy studies.

## References

1. WHO. World Malaria Report. (2022).

2 Saliba, K. J. & Kirk, K. pH regulation in the intracellular malaria parasite, Plasmodium falciparum. H(+) extrusion via a V-type H(+)-ATPase. J Biol Chem 274, 33213–33219 (1999). https://doi.org:10.1074/jbc.274.47.33213

3 Kuhn, Y., Rohrbach, P. & Lanzer, M. Quantitative pH measurements in Plasmodium falciparum-infected erythrocytes using pHluorin. Cell Microbiol 9, 1004–1013 (2007). https://doi.org:10.1111/j.1462-5822.2006.00847.x

4 Bennett, T. N. et al. Drug resistance-associated pfCRT mutations confer decreased Plasmodium falciparum digestive vacuolar pH. Mol Biochem Parasitol 133, 99–114 (2004). https://doi.org:10.1016/j.molbiopara.2003.09.008

5 Saliba, K. J. et al. Acidification of the malaria parasite’s digestive vacuole by a H+-ATPase and a H+-pyrophosphatase. J Biol Chem 278, 5605–5612 (2003). https://doi.org:10.1074/jbc.M208648200

6 Klonis, N. et al. Evaluation of pH during cytostomal endocytosis and vacuolar catabolism of haemoglobin in Plasmodium falciparum. Biochem J 407, 343–354 (2007). https://doi.org:10.1042/BJ20070934

7 Mohring, F. et al. Determination of glutathione redox potential and pH value in subcellular compartments of malaria parasites. Free Radic Biol Med 104, 104–117 (2017). https://doi.org:10.1016/j.freeradbiomed.2017.01.001

8 Casey, J. R., Grinstein, S. & Orlowski, J. Sensors and regulators of intracellular pH. Nat Rev Mol Cell Biol 11, 50–61 (2010). https://doi.org:10.1038/nrm2820

9 Cosse, M. & Seidel, T. Plant Proton Pumps and Cytosolic pH-Homeostasis. Front Plant Sci 12, 672873 (2021). https://doi.org:10.3389/fpls.2021.672873

10 Ginsburg, H. Abundant proton pumping in Plasmodium falciparum, but why? Trends Parasitol 18, 483–486 (2002). https://doi.org:10.1016/s1471-4922(02)02350-4

11 McIntosh, M. T. & Vaidya, A. B. Vacuolar type H+ pumping pyrophosphatases of parasitic protozoa. Int J Parasitol 32, 1–14 (2002). https://doi.org:10.1016/s0020-7519(01)00325-3

12 McIntosh, M. T., Drozdowicz, Y. M., Laroiya, K., Rea, P. A. & Vaidya, A. B. Two classes of plant-like vacuolar-type H(+)-pyrophosphatases in malaria parasites. Mol Biochem Parasitol 114, 183–195 (2001). https://doi.org:10.1016/s0166-6851(01)00251-1

13 Luo, S., Marchesini, N., Moreno, S. N. & Docampo, R. A plant-like vacuolar H(+)-pyrophosphatase in Plasmodium falciparum. FEBS Lett 460, 217–220 (1999). https://doi.org:10.1016/s0014-5793(99)01353-8

14 Karcz, S. R., Herrmann, V. R. & Cowman, A. F. Cloning and characterization of a vacuolar ATPase A subunit homologue from Plasmodium falciparum. Mol Biochem Parasitol 58, 333–344 (1993). https://doi.org:10.1016/0166-6851(93)90056-4

15 Omobukola Solebo, L. L., Jing Zhou, Tian-min Fu, Hangjun Ke. Malaria parasites utilize pyrophosphate to fuel an essential proton pump in the ring stage and the transition to trophozoite stage. BioRxiv (2021). https://doi.org:doi: https://doi.org/10.1101/2021.10.25.465524

16 Vasanthakumar, T. & Rubinstein, J. L. Structure and Roles of V-type ATPases. Trends Biochem Sci 45, 295–307 (2020). https://doi.org:10.1016/j.tibs.2019.12.007

17 Kane, P. M. Disassembly and reassembly of the yeast vacuolar H(+)-ATPase in vivo. J Biol Chem 270, 17025–17032 (1995).

18 Wang, H. & Rubinstein, J. L. CryoEM of V-ATPases: Assembly, disassembly, and inhibition. Curr Opin Struct Biol 80, 102592 (2023). https://doi.org:10.1016/j.sbi.2023.102592

19 Zhang, M. et al. Uncovering the essential genes of the human malaria parasite Plasmodium falciparum by saturation mutagenesis. Science 360 (2018). https://doi.org:10.1126/science.aap7847

20 Bushell, E. et al. Functional Profiling of a Plasmodium Genome Reveals an Abundance of Essential Genes. Cell 170, 260–272 e268 (2017). https://doi.org:10.1016/j.cell.2017.06.030

21 Harrison, M. A. & Muench, S. P. The Vacuolar ATPase - A Nano-scale Motor That Drives Cell Biology. Subcell Biochem 87, 409–459 (2018). https://doi.org:10.1007/978-981-10-7757-9_14

22 Graham, L. A., Flannery, A. R. & Stevens, T. H. Structure and assembly of the yeast V-ATPase. J Bioenerg Biomembr 35, 301–312 (2003). https://doi.org:10.1023/a:1025772730586

23 Lupanga, U. et al. The Arabidopsis V-ATPase is localized to the TGN/EE via a seed plant-specific motif. Elife 9 (2020). https://doi.org:10.7554/eLife.60568

24 Yee, D. P. et al. The V-type ATPase enhances photosynthesis in marine phytoplankton and further links phagocytosis to symbiogenesis. Curr Biol 33, 2541–2547 e2545 (2023). https://doi.org:10.1016/j.cub.2023.05.020

25 Karcz, S. R., Herrmann, V. R., Trottein, F. & Cowman, A. F. Cloning and characterization of the vacuolar ATPase B subunit from Plasmodium falciparum. Mol Biochem Parasitol 65, 123–133 (1994). https://doi.org:10.1016/0166-6851(94)90121-x

26 Hayashi, M. et al. Vacuolar H(+)-ATPase localized in plasma membranes of malaria parasite cells, Plasmodium falciparum, is involved in regional acidification of parasitized erythrocytes. J Biol Chem 275, 34353–34358 (2000). https://doi.org:10.1074/jbc.M003323200

27 Marchesini, N., Vieira, M., Luo, S., Moreno, S. N. & Docampo, R. A malaria parasite-encoded vacuolar H(+)-ATPase is targeted to the host erythrocyte. J Biol Chem 280, 36841–36847 (2005). https://doi.org:10.1074/jbc.M507727200

28 Rajaram, K., Liu, H. B. & Prigge, S. T. Redesigned TetR-Aptamer System To Control Gene Expression in Plasmodium falciparum. mSphere 5 (2020). https://doi.org:10.1128/mSphere.00457-20

29 Ganesan, S. M., Falla, A., Goldfless, S. J., Nasamu, A. S. & Niles, J. C. Synthetic RNA-protein modules integrated with native translation mechanisms to control gene expression in malaria parasites. Nat Commun 7, 10727 (2016). https://doi.org:10.1038/ncomms10727

30 Wagner, J. C., Platt, R. J., Goldfless, S. J., Zhang, F. & Niles, J. C. Efficient CRISPR-Cas9-mediated genome editing in Plasmodium falciparum. Nat Methods 11, 915–918 (2014). https://doi.org:10.1038/nmeth.3063

31 Ghorbal, M. et al. Genome editing in the human malaria parasite Plasmodium falciparum using the CRISPR-Cas9 system. Nat Biotechnol 32, 819–821 (2014). https://doi.org:10.1038/nbt.2925

32 Ho, C. M. et al. Malaria parasite translocon structure and mechanism of effector export. Nature 561, 70–75 (2018). https://doi.org:10.1038/s41586-018-0469-4

33 Ressurreicao, M. et al. Use of a highly specific kinase inhibitor for rapid, simple and precise synchronization of Plasmodium falciparum and Plasmodium knowlesi asexual blood-stage parasites. PLoS One 15, e0235798 (2020). https://doi.org:10.1371/journal.pone.0235798

34 Krugliak, M., Zhang, J. & Ginsburg, H. Intraerythrocytic Plasmodium falciparum utilizes only a fraction of the amino acids derived from the digestion of host cell cytosol for the biosynthesis of its proteins. Mol Biochem Parasitol 119, 249–256 (2002). https://doi.org:10.1016/s0166-6851(01)00427-3

35 Goldberg, D. E. Hemoglobin degradation in Plasmodium-infected red blood cells. Semin Cell Biol 4, 355–361 (1993). https://doi.org:10.1006/scel.1993.1042

36 Spielmann, T., Gras, S., Sabitzki, R. & Meissner, M. Endocytosis in Plasmodium and Toxoplasma Parasites. Trends Parasitol 36, 520–532 (2020). https://doi.org:10.1016/j.pt.2020.03.010

37 Jonscher, E. et al. PfVPS45 Is Required for Host Cell Cytosol Uptake by Malaria Blood Stage Parasites. Cell Host Microbe 25, 166–173 e165 (2019). https://doi.org:10.1016/j.chom.2018.11.010

38 Milani, K. J., Schneider, T. G. & Taraschi, T. F. Defining the morphology and mechanism of the hemoglobin transport pathway in Plasmodium falciparum-infected erythrocytes. Eukaryot Cell 14, 415–426 (2015). https://doi.org:10.1128/EC.00267-14

39 Wang, R. et al. Molecular basis of V-ATPase inhibition by bafilomycin A1. Nature Communications 12, 1782 (2021). https://doi.org:10.1038/s41467-021-22111-5

40 Parra, K. J. & Kane, P. M. Reversible association between the V1 and V0 domains of yeast vacuolar H+-ATPase is an unconventional glucose-induced effect. Mol Cell Biol 18, 7064–7074 (1998). https://doi.org:10.1128/MCB.18.12.7064

41 Parra, K. J., Keenan, K. L. & Kane, P. M. The H subunit (Vma13p) of the yeast V-ATPase inhibits the ATPase activity of cytosolic V1 complexes. J Biol Chem 275, 21761–21767 (2000). https://doi.org:10.1074/jbc.M002305200

42 Stasic, A. J. et al. The Toxoplasma Vacuolar H(+)-ATPase Regulates Intracellular pH and Impacts the Maturation of Essential Secretory Proteins. Cell Rep 27, 2132–2146 e2137 (2019). https://doi.org:10.1016/j.celrep.2019.04.038

43 Francis, S. E., Sullivan, D. J., Jr. & Goldberg, D. E. Hemoglobin metabolism in the malaria parasite Plasmodium falciparum. Annu Rev Microbiol 51, 97–123 (1997). https://doi.org:10.1146/annurev.micro.51.1.97

44 Liu, J., Istvan, E. S., Gluzman, I. Y., Gross, J. & Goldberg, D. E. Plasmodium falciparum ensures its amino acid supply with multiple acquisition pathways and redundant proteolytic enzyme systems. Proc Natl Acad Sci U S A 103, 8840–8845 (2006). https://doi.org:10.1073/pnas.0601876103

45 Ludwig, J. et al. Identification and characterization of a novel 9.2-kDa membrane sector-associated protein of vacuolar proton-ATPase from chromaffin granules. J Biol Chem 273, 10939–10947 (1998). https://doi.org:10.1074/jbc.273.18.10939

46 Alder, A. et al. The role of Plasmodium V-ATPase in vacuolar physiology and antimalarial drug uptake. Proc Natl Acad Sci U S A 120, e2306420120 (2023). https://doi.org:10.1073/pnas.2306420120

47 Hameed, P. S. et al. Triaminopyrimidine is a fast-killing and long-acting antimalarial clinical candidate. Nat Commun 6, 6715 (2015). https://doi.org:10.1038/ncomms7715

48 Reader, J. et al. Multistage and transmission-blocking targeted antimalarials discovered from the open-source MMV Pandemic Response Box. Nat Commun 12, 269 (2021). https://doi.org:10.1038/s41467-020-20629-8

49 Meibalan, E. et al. Host erythrocyte environment influences the localization of exported protein 2, an essential component of the Plasmodium translocon. Eukaryot Cell 14, 371–384 (2015). https://doi.org:10.1128/EC.00228-14

50 Ling, L. et al. Genetic ablation of the mitoribosome in the malaria parasite Plasmodium falciparum sensitizes it to antimalarials that target mitochondrial functions. J Biol Chem 295, 7235–7248 (2020). https://doi.org:10.1074/jbc.RA120.012646

51 Ke, H., Dass, S., Morrisey, J. M., Mather, M. W. & Vaidya, A. B. The mitochondrial ribosomal protein L13 is critical for the structural and functional integrity of the mitochondrion in Plasmodium falciparum. J Biol Chem 293, 8128–8137 (2018). https://doi.org:10.1074/jbc.RA118.002552

